# Subcortical Anatomy of the Default Mode Network: a functional and structural connectivity study

**DOI:** 10.1101/528679

**Authors:** Pedro Nascimento Alves, Chris Foulon, Vyacheslav Karolis, Danilo Bzdok, Daniel S. Margulies, Emmanuelle Volle, Michel Thiebaut de Schotten

**Affiliations:** Brain Connectivity and Behaviour Laboratory, BCBlab, Sorbonne Universities, Paris France.; Frontlab, Institut du Cerveau et de la Moelle épinière (ICM), UPMC UMRS 1127, Inserm U 1127, CNRS UMR 7225, Paris, France.; Department of Neurosciences and Mental Health, Neurology, Hospital de Santa Maria, CHLN, Lisbon, Portugal.; Language Research Laboratory, Faculty of Medicine, Universidade de Lisboa, Lisbon, Portugal.; Computational Neuroimaging Laboratory, Department of Diagnostic Medicine, The University of Texas at Austin Dell Medical School; INRIA, Parietal Team, Saclay, France; Neurospin, CEA, Gif-sur-Yvette, France; Department of Psychiatry, Psychotherapy and Psychosomatics, RWTH Aachen University, Aachen, Germany; JARA-BRAIN, Jülich-Aachen Research Alliance, Germany; Centre de Neuroimagerie de Recherche CENIR, Groupe Hospitalier Pitié-Salpêtrière, Paris, France.; Groupe d’Imagerie Neurofonctionnelle, Institut des Maladies Neurodégénératives-UMR 5293, CNRS, CEA University of Bordeaux, Bordeaux, France

## Abstract

Most existing research into the default-mode network (DMN) has taken a corticocentric approach. Despite the resemblance of the DMN with the unitary model of the limbic system, the anatomy and contribution of subcortical structures to the network may be underappreciated due to methods limitation. Here, we propose a new and more comprehensive neuroanatomical model of the DMN including the basal forebrain and anterior and mediodorsal thalamic nuclei and cholinergic nuclei. This has been achieved by considering functional territories during interindividual brain alignment. Additionally, tractography of diffusion-weighted imaging was employed to explore the structural connectivity of the DMN and revealed that the thalamus and basal forebrain had high importance in term of values of node degree and centrality in the network. The contribution of these neurochemically diverse brain nuclei reconciles previous neuroimaging with neuropathological findings in diseased brain and offers the potential for identifying a conserved homologue of the DMN in other mammalian species.

## 1. Introduction

For the first time in 1979, David Ingvar used Xenon clearance to investigate resting wakefulness (Ingvar, 1979). When aligned by scalp and skull markers, the 11 brains examined indicated an evident increase of the blood flow levels in the frontal lobe interpreted as a surrogate for undirected, spontaneous, conscious mental activity. Later, Positron Emission Tomography (PET) was used to map more systematically task-related activation in the brain, often with resting wakefulness as a control task. The contrast between task-related and resting wakefulness led to the description of deactivation (i.e. active at rest more than during the task) in a set of regions including retrosplenial cortex, inferior parietal cortex, dorsolateral frontal cortex inferior frontal cortex, left inferior temporal gyrus, medial frontal regions and amygdala (Mazoyer et al., 2001; Shulman et al., 1997a) that quickly bore the name of default mode network (DMN) (Raichle et al., 2001). In these studies, skull landmarks or structural Magnetic Resonance Imaging (MRI) were used to align PET images in Talairach stereotaxic or in Montreal Neurological Institute (MNI) templates (Mazoyer et al., 2001; Shulman et al., 1997a). The advent of functional Magnetic Resonance Imaging (fMRI), particularly of methods for analysing functional connectivity, led to the allocation of new structures to this network, such as the hippocampal formation (Buckner et al., 2008; Greicius et al., 2004; Vincent et al., 2006).

Today, the DMN has largely been a cortically-defined set of network nodes. Consisting of distinct regions/nodes distributed across the ventromedial and lateral prefrontal, posteromedial and inferior parietal, as well as lateral and medial temporal cortex, the DMN is considered a backbone of cortical integration (Andrews-Hanna et al., 2010; Kernbach et al., 2018; Lopez-Persem et al., 2018; Margulies et al., 2016). Its subcortical components are, however, less well characterized. Studies of whole-brain network organization reveal subregions of the cerebellum (Buckner et al., 2011; Stoodley and Schmahmann, 2009) and striatum (Choi et al., 2012) that are functionally connected with the cortical regions of the DMN. Seed-based functional connectivity studies further demonstrate additional DMN-specific connectivity to several subcortical structures, including the amygdala (Bzdok et al., 2012; Roy et al., 2009), striatum (Di Martino et al., 2008), thalamus (Fransson, 2005). These studies are important as a cleaner characterization of the anatomy of the DMN is an essential step towards understanding its functional role and its involvement in brain diseases. Particularly, an increased activity characterises the regions that compose DMN during tasks involving autobiographical, episodic and semantic memory, mind wandering, perspective-taking or future thinking (Shapira-Lichter et al., 2013; Shulman et al., 1997b; Bendetowicz et al. 2018). Conversely, DMN regions show a decreased neural activity during attention-demanding and externally-oriented tasks (Shulman et al., 1997b; Spreng et al., 2009). Finally, altered connectivity in the DMN has been observed in a large variety of brain diseases, including Alzheimer’s disease, Parkinson’s disease, schizophrenia, depression, temporal lobe epilepsy, attention deficit and hyperactivity disorder, drug addiction, among others (Broyd et al., 2009; Geng et al., 2017; Tessitore et al., 2012; Voets et al., 2012; Zhu et al., 2017). Hence, while prior research provides first hints towards a broader definition of the DMN system, further research is necessary to articulate the anatomical extent of specific subcortical contributions, and to understand the independent contribution of these structures in DMN function and pathologies.

Nevertheless, since the DMN has repeatedly been characterized as a cohesive functional network (Buckner et al., 2008), an average of brain images relying exclusively on anatomical references and landmarks may be suboptimal (Brett et al., 2002; Thiebaut de Schotten and Shallice, 2017) whether the method employed is a surface-based or volume-based registration (Brett et al., 2002; Despotovic et al., 2015). Small structures of the brain may be particularly susceptible to this misalignment, especially when MRI lacks contrast. Besides morphology, cytoarchitecture and function are poorly overlapping, especially in the DMN(Bzdok et al., 2015; Eickhoff et al., 2016). Consequently, functional areas present in every subject may not overlap after averaging all structurally aligned brain images in a group analysis (Braga and Buckner, 2017; Brett et al., 2002). This biological misalignment can be particularly problematic for revealing significant small regions of the DMN (figure 1). A better alignment is also essential for the subcortical structures of the brain as their variability is still considerable (Amunts et al., 2005, 1999; Croxson et al., 2017; Zaborszky et al., 2008). Specifically, cytoarchitectonic studies have shown that only one-quarter of the volume of cholinergic nuclei overlaps in at least half of the individuals studied (Zaborszky et al., 2008). Similarly, structures such as mammillary bodies, nucleus basalis of Meynert, or anterior thalamic nuclei, can vary in size, morphology and locations, and are particularly prone to misalignment with the current methods of structural registration (Despotovic et al., 2015; Liu et al., 2015; Möttönen et al., 2015; Tagliamonte et al., 2013). To address several of these challenges, we propose to revisit the anatomical scaffold of the DMN using a coregistration based on a functional alignment, rather than on exclusively structural landmarks. This approach has already been used to overcome the high interindividual variability of the morphology of some areas of heteromodal association cortex and led to a more accurate mapping of resting-state functional connectivity (Langs et al., 2015; Mueller et al., 2013).

**Figure 1:**
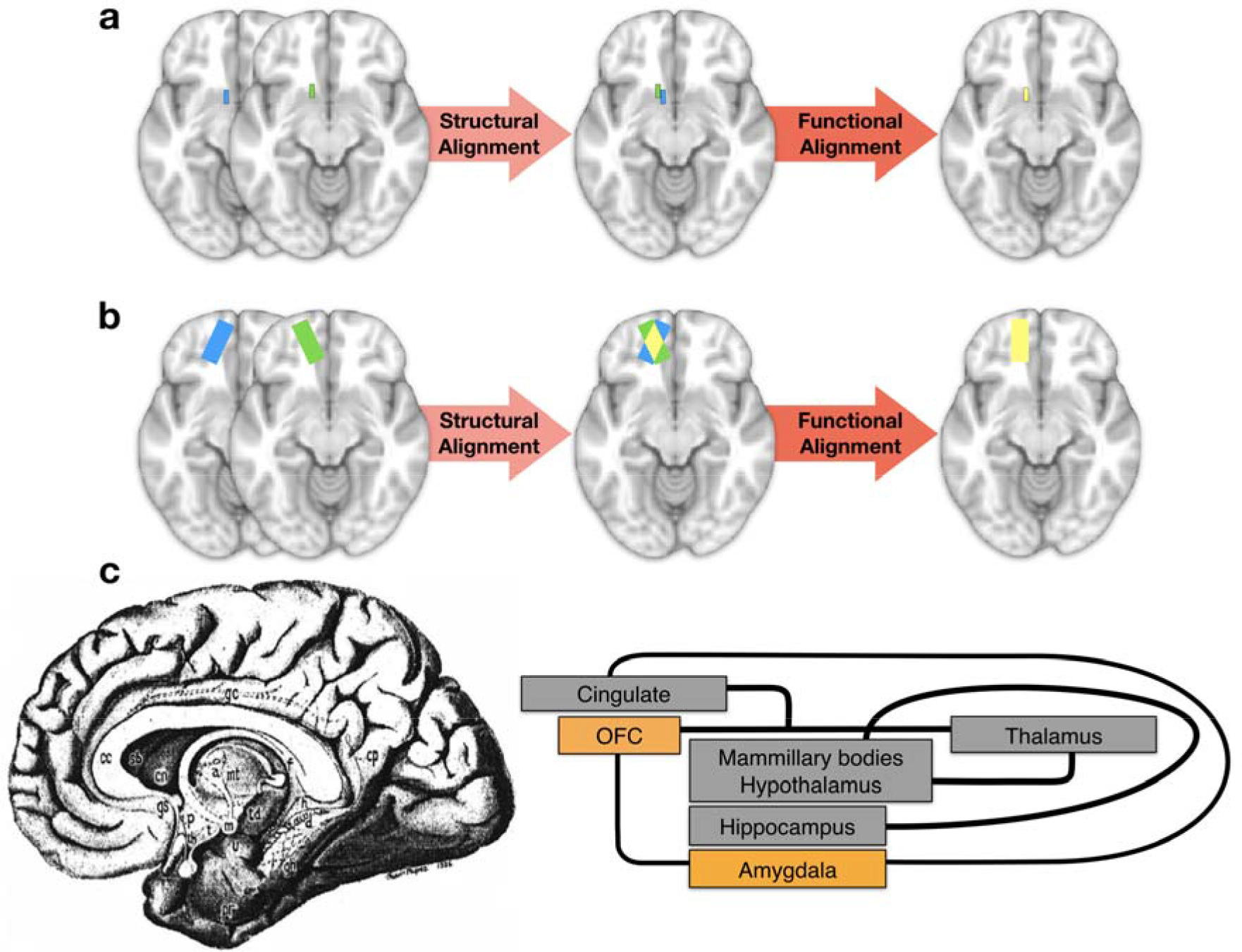
Illustration of intersubject alignment of brain images & unitary model of the limbic system. Blue and green rectangles represent the same functional area in two subjects, while yellow rectangles illustrate the overlap of the two individual areas after alignment. With structural alignment, there can be a complete misalignment of small functional areas (a) or partial misalignment of large functional areas (b) due to functional-anatomical variability or poor anatomical contrasts in MRI imaging. If an additional step of functional alignment is performed, an optimized overlap of functional areas is obtained. (c) The limbic system (left panel) as originally depicted by Papez in (1937) and (right panel) diagram of the unitary model of the limbic system (MacLean, 1952, 1949; Catani et al. 2013)

We hypothesised that using a functional alignment will reveal structures of basal forebrain and the Papez’s circuits, namely anterior and mediodorsal thalamic nuclei and mammillary bodies, as constituent nodes of the DMN for several reasons. First, all these regions are highly interconnected which suggest they belong to the same functional system (Yakovlev, 1948; Yakovlev and Locke, 1961). Second, the current conceptualization of DMN anatomy resembles the unitary model of the limbic system (**Figure 1c**) which, through the coordination of its subregions, subserves the elaboration of emotion, memories and behaviour (Catani et al., 2013; MacLean, 1952, 1949; Papez, 1937). Third, the basal forebrain comprises a group of neurochemically diverse nuclei, involved in dopaminergic, cholinergic and serotoninergic pathways, that are crucial in the pathophysiology of the aforementioned diseases that affect the DMN connectivity. Finally, recent electrophysiological evidence has shown that in rats the basal forebrain exhibits the same pattern of gamma oscillations than DMN and that it influences the activity of anterior cingulate cortex (Nair et al., 2018).

In this study, we used a functional alignment of rs-*f*MRI-based individual DMN maps to build a more comprehensive DMN model. To provide a complete window into the anatomy of the DMN, we explored the structural connectivity of our new model of the DMN using tractography imaging techniques. We combined the tractography results with graph measures to corroborate the essential contribution of the new regions reported to the DMN.

## 2. Material and Methods

### 2.1. Subjects and MRI acquisition

MRI images of subjects without neurological or psychiatric disease were obtained (age mean±SD 29±6 years, range 22-42 years; 11 female, 9 male) with a Siemens 3 Tesla Prisma system equipped with a 64-channel head coil.

An axial 3D T1-weighted imaging dataset covering the whole head was acquired for each participant (286 slices, voxel resolution = 0.7 mm3, echo time (TE) = 2.17 ms, repetition time (TR) = 2400 ms, flip angle = 9°).

Resting state functional MRI (rs-*f*MRI) images were obtained using T2*-weighted echo-planar imaging (EPI) with blood oxygenation level-dependent (BOLD) contrast. EPIs (TR/TE = 2050/25 ms) comprised 42 axial slices acquired with a multiband pulse (Feinberg et al., 2010; Moeller et al., 2010; Setsompop et al., 2012; Xu et al., 2013) covering the entire cerebrum (voxel size = 3 mm3) including 290 brain scan volumes in one run of 10 minutes.

A diffusion-weighted imaging (DWI) acquisition sequence, fully optimised for tractography, provided isotropic (1.7 × 1.7 × 1.7 mm) resolution and coverage of the whole head with a posterior-anterior phase of acquisition, with an echo time (TE) = 75 msec. A repetition time (TR) equivalent to 3500ms was used. At each slice location, six images were acquired with no diffusion gradient applied (b-value of 0 sec mm−2). Additionally, 60 diffusion-weighted images were acquired, in which gradient directions were uniformly distributed on the hemisphere with electrostatic repulsion. The diffusion weighting was equal to a b-value of 2000 s/mm−2. This sequence was fully repeated with reversed phase-encode blips. This provides us with two datasets with distortions going in opposite directions. From these pairs, the susceptibility-induced off-resonance field was estimated using a method similar to that described in (Andersson et al., 2003) and corrected on the whole diffusion-weighted dataset using the tool TOPUP and EDDY as implemented in FSL (Smith et al., 2004).

### 2.2. Resting-state functional MRI analysis - overview

Resting state functional MRI images were corrected for artefacts in Funcon-Preprocessing tool of the Brain Connectivity and Behaviour toolkit (http://toolkit.bcblab.com; Foulon et al., 2018; Jenkinson et al., 2002; Woolrich et al., 2009). In a next step, they were registered to T1 high resolution individual structural images and normalized to MNI152 standard space using Advanced Normalization Tools (ANTS; http://stnava.github.io/ANTs; Avants and Gee, 2004; Avants et al., 2010).

#### 2.2.1 Individual DMN maps in the structural space

Individual subject-tailored/fitted DMN maps were obtained by correlation with seed regions of interest of a functional parcellated brain template. The regions used for the seed-based functional connectivity analysis were those defined as DMN regions in the resting-state parcellation map by Gordon and collaborators (Gordon et al., 2016). Gordon and collaborators created these parcellations according to abrupt changes in resting state’s time course profile, each parcel having a homogenous time course profile. This general-purpose atlas provides a total of 40 DMN nodes (20 in each hemisphere), which were used as seeds for building the correlation maps (http://toolkit.bcblab.com; Foulon et al., 2018). This produced a set of 40 maps per subjects.

The individual DMN map was obtained by calculating the median of the 40 seed-based correlation maps of each individual using FSL (Jenkinson et al., 2012).

To obtain a group DMN map in the structural space, the median of the twenty individual maps was derived.

#### 2.2.2. Individual DMN maps in the “functional space”

To achieve the proposed optimized map of the DMN, the same twenty individual maps were functionally aligned with each other in a new “functional space” using the following steps: Individual DMN maps (obtained in 2.2.1.) were aligned with each other using ANTs' script buildtemplateparallel.sh, defining cross-correlation as the similarity measure and greedy SyN as the transformation model (Avants et al., 2011, 2008; Klein et al., 2009). This approach consists in an iterative (n=4) diffeomorphic transformation to a common space. The group map was obtained by calculating the median of all DMN maps after functional alignment. The resulting maps correspond to alignment of the 20 individual DMN maps in a “functional space” (figure 2).

**Figure 2:**
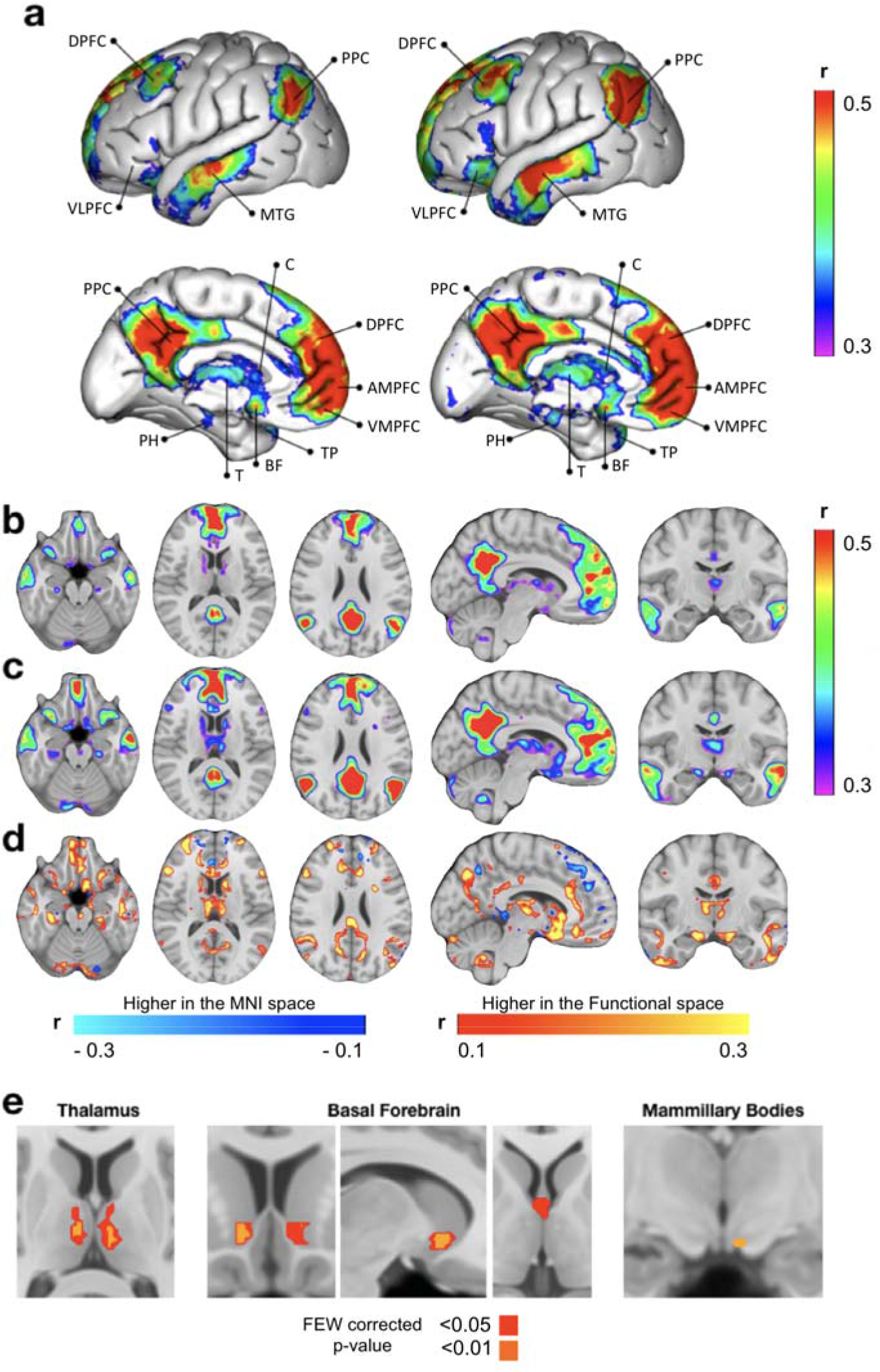
Maps of the DMN structurally or functionally aligned. (a) 3D view of the two DMN left panel corresponds to the structural space alignment, right panel to the functional space alignment, (b) brain sections of the structurally aligned DMN, (c) brain sections of the functionally aligned DMN, (d) subtraction of the structurally and the functionally aligned DMN maps, (e) paired t-test between the structurally and the functionally aligned DMN in the three hypothesized regions, with colours indicating statistically significant differences at two levels of significance - <0.05 and <0.01, family-wise error (FWE) corrected p-values (higher in the functional space). DPFC, Dorsal Prefrontal Cortex; PPC, Posterior Parietal Cortex; VLPFC, Ventro-Lateral Prefrontal Cortex; MTG, Middle Temporal Gyrus; PCC, Posterior Cingulate Cortex; C, Caudate; DPFC, Dorsal Prefrontal Cortex; AMPFC, Antero-Median Prefrontal Cortex; VMPFC, Ventro-Median Prefrontal Cortex; TP, Temporal Pole; BF, Basal Forebrain; T, Thalamus; PH, Parahippocampal.

#### 2.2.3. Functional connectivity comparison of the two DMN

For the functional and structural-based DMN maps, time-series of rs-*f*MRI of each individual in the different regions of interest identified in the DMN maps were extracted (Jenkinson et al., 2012). Correlation and partial correlation coefficients were calculated as measures of functional connectivity.

Two matrices, representing the median correlation values in the MNI152 space and in the functional space, were created using BrainNet Viewer (Xia et al., 2013). A paired t-test was calculated for each cell of the two connectivity matrices using Python’s Scipy package, version 0.19.1 (https://www.scipy.org/; scipy.stats.ttest_rel()). An uncorrected p-value of <0.05 and a p-value corrected for multiple comparisons at a threshold of <0.0001 (Bonferroni correction) were used. Circos software was used to illustrate functional connections in the functional space (http://circos.ca/; Krzywinski, 2009).

#### 2.2.4. Anatomical validation in thalamic, basal forebrain and mesencephalic areas

Meynert nuclei, medial septal nuclei and diagonal band of Broca probabilistic maps were derived from the work of Zaborszky et al., the Harvard-Oxford probabilistic atlas was employed for nucleus accumbens, Talairach atlas registered to the MNI152 space for mammillary bodies and thalamic nuclei, and Harvard Ascending Arousal Network Atlas for ventral tegmental area (Desikan et al., 2006; Edlow et al., 2012; Lancaster et al., 2007; Talairach and Tournoux, 1988; Zaborszky et al., 2008). A percentage of volume overlap between the DMN map and each nucleus of interest was subsequently calculated for each subject.

### 2.3. Tractography analysis

Diffusion Weighted Images were corrected for signal drift (Vos et al., 2017), motion and eddy current artefacts using ExploreDTI (http://www.exploredti.comd; Leemans et al., 2009).

Whole-brain tractography was performed on the software StarTrack using a deterministic approach (https://www.mr-startrack.com). A damped Richardson-Lucy algorithm was applied for spherical deconvolutions (Dell’Acqua et al., 2010). A fixed fibre response corresponding to a shape factor of α = 1.5×10−3 mm2/s was adopted. The defined number of iterations was 150 and the geometric damping parameter was 8. The absolute threshold was defined as 3 times the spherical fibre orientation distribution (FOD) of a grey matter isotropic voxel and the relative threshold as 8% of the maximum amplitude of the FOD (Thiebaut de Schotten et al., 2014). A modified Euler algorithm was used (Dell’Acqua et al., 2013). An angle threshold of 35°, a step size of 0.85mm, and a minimum length of 20mm were chosen.

Diffusion tensor images were registered to the MNI152 standard space and, then, into the “functional space” applying the affine and diffeomorphic deformation generated in 2.2 and in 2.2.2., using the tool tractmath as part of the software package Tract Querier (Wassermann et al., 2016).

Regions of interest were defined according to the previous anatomical models of the DMN (Andrews-Hanna et al., 2010; Buckner et al., 2008). Additional regions of interest were defined in the novel subcortical regions identified in our new model.

The command tckedit of MRtrix toolbox (http://www.mrtrix.org/; Tournier et al., 2012) was used to isolate the tracts of interest. The map of the DMN created through functional alignment was used to define the regions of interest.

In order to have a group-representative tract volume for each tract, one sample T-test of individual tract volumes was calculated using FSL randomise, with variance smoothing of 4mm. BrainVisa was used to create the corresponding illustration of the tracts that reached significance (Rivière et al., 2011).

#### 2.3.1. Graph theory analysis of structural connectivity

To explore whether the new putative regions of the DMN are essential structures and potential areas of vulnerability in the network, we investigated the hub properties of the network nodes using graph theory measures, namely node degree and betweenness centrality (Bullmore and Sporns, 2009). Node degree refers to the number of connections between a given node and the other nodes of the network. Betweenness centrality is the fraction of all shortest paths in the network that pass through a given node (Bullmore and Sporns, 2009).

For each participant, an anatomical connectome matrix of the DMN was built using the tck2connectome command of MRtrix (http://www.mrtrix.org/; Tournier et al., 2012). Each region of interest was defined as a node. Only the streamlines that ended in both regions of interest were considered (Shu et al., 2011). The matrices were binarised (no threshold) for structural connection or no structural connection between regions. Brain Connectivity Toolbox for Python (https://pypi.python.org/pypi/bctpy) was used to obtain the network measures. The functions degrees_und and betweenness_bin were run, respectively, for each individual matrix (Rubinov and Sporns, 2010). The median values for each measure were obtained. The illustration of the network was made using Surf Ice (https://www.nitrc.org/projects/surfice/)

## 3. Results

### 3.1 Anatomical comparison between the two alignment methods

DMN connectivity maps obtained from structural and functional alignments are displayed in figure 2a-c. In both maps, classical areas of the DMN were observed, namely: posterior cingulate cortex and retrosplenial cortex; ventromedial, anteromedial, and dorsal prefrontal cortex; temporal pole; middle temporal gyrus; hippocampus and parahippocampal cortex; amygdala; and the posterior parietal cortex.

Figure 2d illustrate the simple voxel-based subtraction between the DMN connectivity maps obtained from structural and functional alignments. Higher average connectivity was achieved in the functionally aligned DMN map in large areas, such as medial prefrontal cortex and posterior cingulate cortex, mostly in the border zones. In fact, the highest differences in connectivity between structural and functional aligned DMN were at the level of the basal forebrain and thalamus. These areas were poorly or even not represented with alignment in the MNI152 space, but were visible after the functional alignment. DMN connectivity was also visible in the medial mesencephalic region, as well in inferior regions in the caudate nuclei, ventrolateral prefrontal cortex, cerebellar tonsils and cerebellar hemispheres. As expected, a significant difference was found between the two maps bilaterally in the thalamus and in the basal forebrain, and in a peripheral zone of the left mammillary body (figure 2d, Table 1). Unthresholded statistical maps are available at NeuroVault.org (https://neurovault.org/collections/OCAMCQFK/).

**Table 1.**
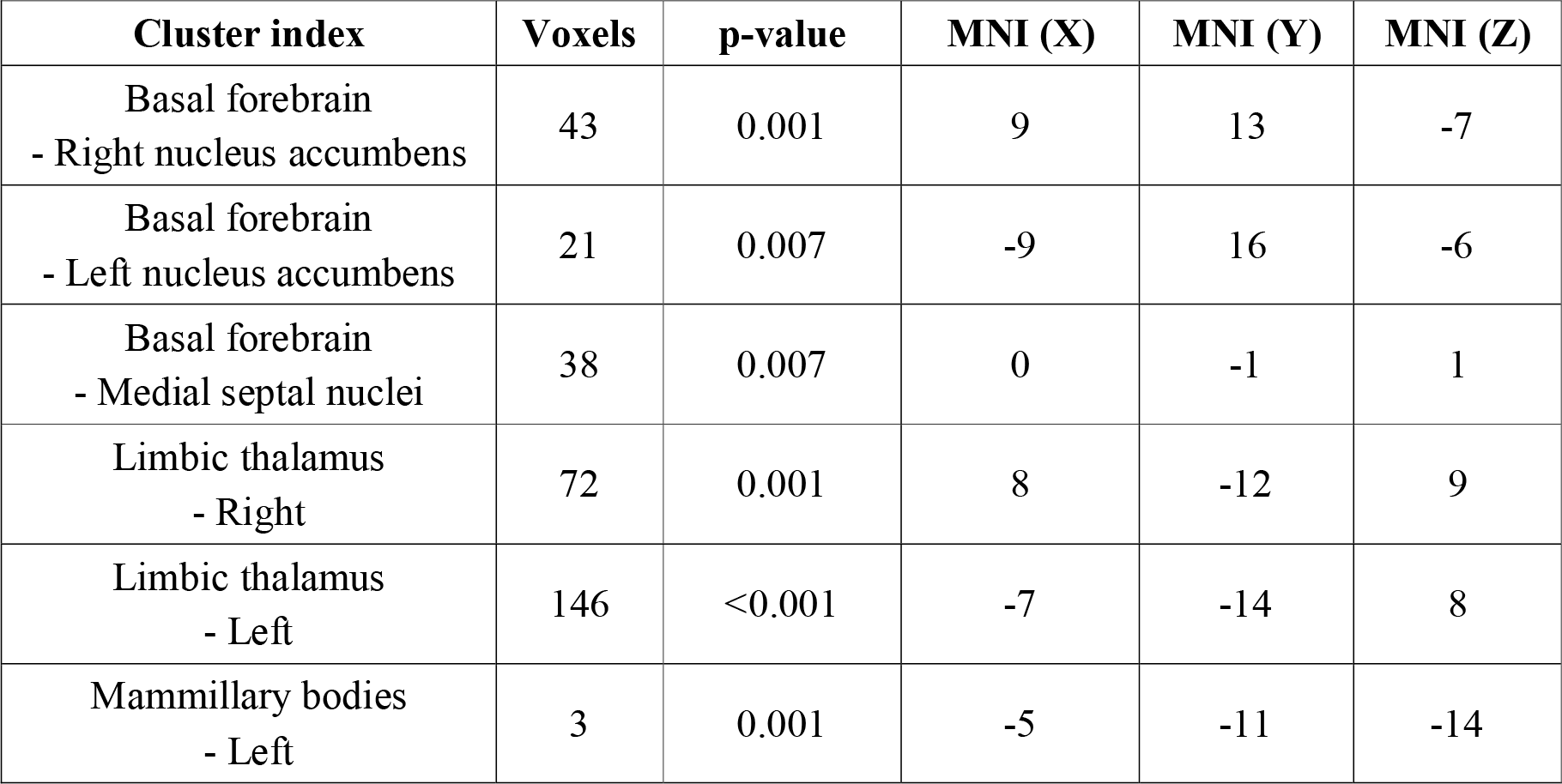
Clusters of the statistical maps obtained when comparing the two methods of alignment. Coordinates represent the centre of gravity and ‘p-value’ of the lowest value found in the cluster.

| **Cluster index** | **Voxels** | **p-value** | **MNI (X)** | **MNI (Y)** | **MNI (Z)** |
| --- | --- | --- | --- | --- | --- |
| Basal forebrain<br>- Right nucleus accumbens | 43 | 0.001 | 9 | 13 | -7 |
| Basal forebrain<br>- Left nucleus accumbens | 21 | 0.007 | -9 | 16 | -6 |
| Basal forebrain<br>- Medial septal nuclei | 38 | 0.007 | 0 | -1 | 1 |
| Limbic thalamus<br>- Right | 72 | 0.001 | 8 | -12 | 9 |
| Limbic thalamus<br>- Left | 146 | <0.001 | -7 | -14 | 8 |
| Mammillary bodies<br>- Left | 3 | 0.001 | -5 | -11 | -14 |

### 3.2 Functional connectivity of the DMN in the functional space

The association strength determined by Pearson’s correlation between the rs-*f*MRI time-series of the regions of interest (i.e. functional connectivity) were higher with alignment in the functional space, compared with structural space, in all pairs of regions (figure 3a). The difference was statistically significant in 95% and in 18% of pairs, whether or not correcting and correcting for multiple comparisons was applied (figure 3b). Table 2 represent the MNI coordinates of all the regions of interest in the DMN map.

**Table 2.** Regions of interest. MNI coordinates, represent the centre of gravity of each region.

| **Region of interest** | **Voxels** | **MNI (X)** | **MNI (Y)** | **MNI (Z)** |
| --- | --- | --- | --- | --- |
| Left Ventro-Median Prefrontal Cortex | 2267 | -11 | 55 | -5 |
| Right Ventro-Median Prefrontal Cortex | 2673 | 11 | 53 | -6 |
| Left Antero-Median Prefrontal Cortex | 2243 | -10 | 50 | 20 |
| Right Antero-Median Prefrontal Cortex | 2144 | 10 | 50 | 19 |
| Left Dorsal Prefrontal Cortex | 3818 | -20 | 31 | 46 |
| Right Dorsal Prefrontal Cortex | 3084 | 23 | 32 | 46 |
| Left Posterior Cingulate Cortex | 2484 | -5 | -50 | 35 |
| Right Posterior Cingulate Cortex | 2224 | 7 | -51 | 34 |
| Left Retrosplenial Cortex | 845 | -6 | -55 | 12 |
| Right Retrosplenial Cortex | 638 | 6 | -54 | 13 |
| Left Posterior Parietal Cortex | 2448 | -46 | -64 | 33 |
| Right Posterior Parietal Cortex | 1733 | 50 | -59 | 34 |
| Left Middle Temporal Gyrus | 2406 | -58 | -21 | -15 |
| Right Middle Temporal Gyrus | 2170 | 59 | -17 | -18 |
| Left Temporal Pole | 348 | -38 | 17 | -34 |
| Right Temporal Pole | 318 | 43 | 15 | -35 |
| Left Ventro-Lateral Cortex | 706 | -36 | 23 | -16 |
| Right Ventro-Lateral Cortex | 487 | 37 | 25 | -16 |
| Left Parahippocampal Region | 355 | -24 | -30 | -16 |
| Right Parahippocampal Region | 287 | 26 | -26 | -18 |
| Left Amygdala | 66 | -15 | -9 | -18 |
| Right Amygdala | 58 | 17 | -8 | -16 |
| Left Caudate | 303 | -11 | 12 | 7 |
| Right Caudate | 266 | 13 | 11 | 9 |
| Left Cerebellar Hemisphere | 906 | -26 | -82 | -33 |
| Right Cerebellar Hemisphere | 1500 | 29 | -79 | -34 |
| Left Cerebellar Tonsil | 184 | -6 | -57 | -45 |
| Right Cerebellar Tonsil | 278 | 8 | -53 | -48 |
| Left Thalamus | 382 | -7 | -14 | 8 |
| Right Thalamus | 305 | 7 | -11 | 8 |
| Left Basal Forebrain | 456 | -7 | 12 | -12 |
| Right Basal Forebrain | 351 | 7 | 9 | -12 |
| Midbrain | 65 | -1 | -22 | -21 |

**Figure 3:**
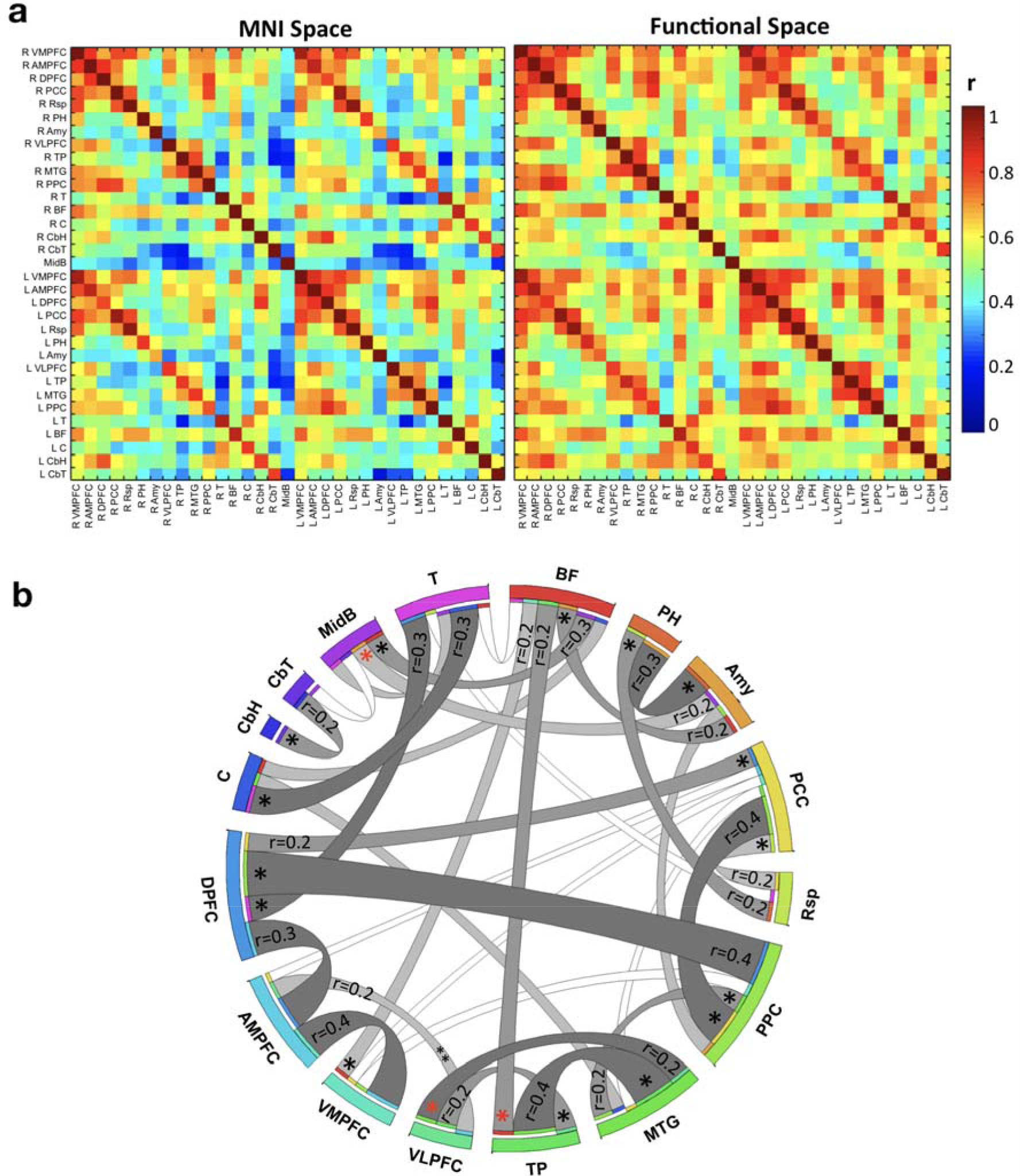
Functional connectivity. a) matrices of the Pearson’s correlations between rs-*f*MRI time-series of the regions of interest in the structural and in the functional space b) graph representation of the functional connectivity between regions of interest in the functional space (connections with partial correlation above 0.2). The level of significance of the difference between Pearson’s correlation in the structural and in functional spaces is indicated as follows: black asterisk p<0.05; red asterisk p<0.0001 (p-value corrected for multiple comparisons). The left side structures are not represented, for a clearer visualization. DPFC, Dorsal Prefrontal Cortex; PPC, Posterior Parietal Cortex; VLPFC, Ventro-Lateral Prefrontal Cortex; Rsp, Retrosplenial Cortex MTG, Middle Temporal Gyrus; PCC, Posterior Cingulate Cortex; C, Caudate; DPFC, Dorsal Prefrontal Cortex; AMPFC, Antero-Median Prefrontal Cortex; VMPFC, Ventro-Median Prefrontal Cortex; TP, Temporal Pole; BF, Basal Forebrain; T, Thalamus; PH, Parahippocampal Region; CbH, Cerebellar Hemisphere; CbT, Cerebellar Tonsil; Amy, Amygdala; MidB, Midbrain.

Regarding the hypothesized areas, basal forebrain had stronger partial correlations with midbrain, ventromedial prefrontal cortex, amygdala and temporal pole, while thalamus had higher partial correlations with caudate nucleus and dorsal prefrontal cortex (figure 3b).

### 3.3 Anatomical validation in thalamic, basal forebrain and mesencephalic areas

Figure 4 illustrates the intersection of the new DMN map after its translation to the MNI group space using individual inverse transformation matrices specific for each individual. All subjects’ DMN spatially overlapped with the templates of the left anterior thalamic nucleus, mediodorsal thalamic nuclei, medial septal nuclei and left nucleus accumbens (Table 3) (Desikan et al., 2006; Edlow et al., 2012; Lancaster et al., 2007; Talairach and Tournoux, 1988; Zaborszky et al., 2008). The number of subjects with an intersection with the right anterior thalamic nucleus, right nucleus accumbens and ventral tegmental area was also very high (95%, 95% and 90% respectively), while the intersection with the other basal forebrain nuclei occurred in approximately half of the subjects, possibly due to their very small size.

**Table 3.**
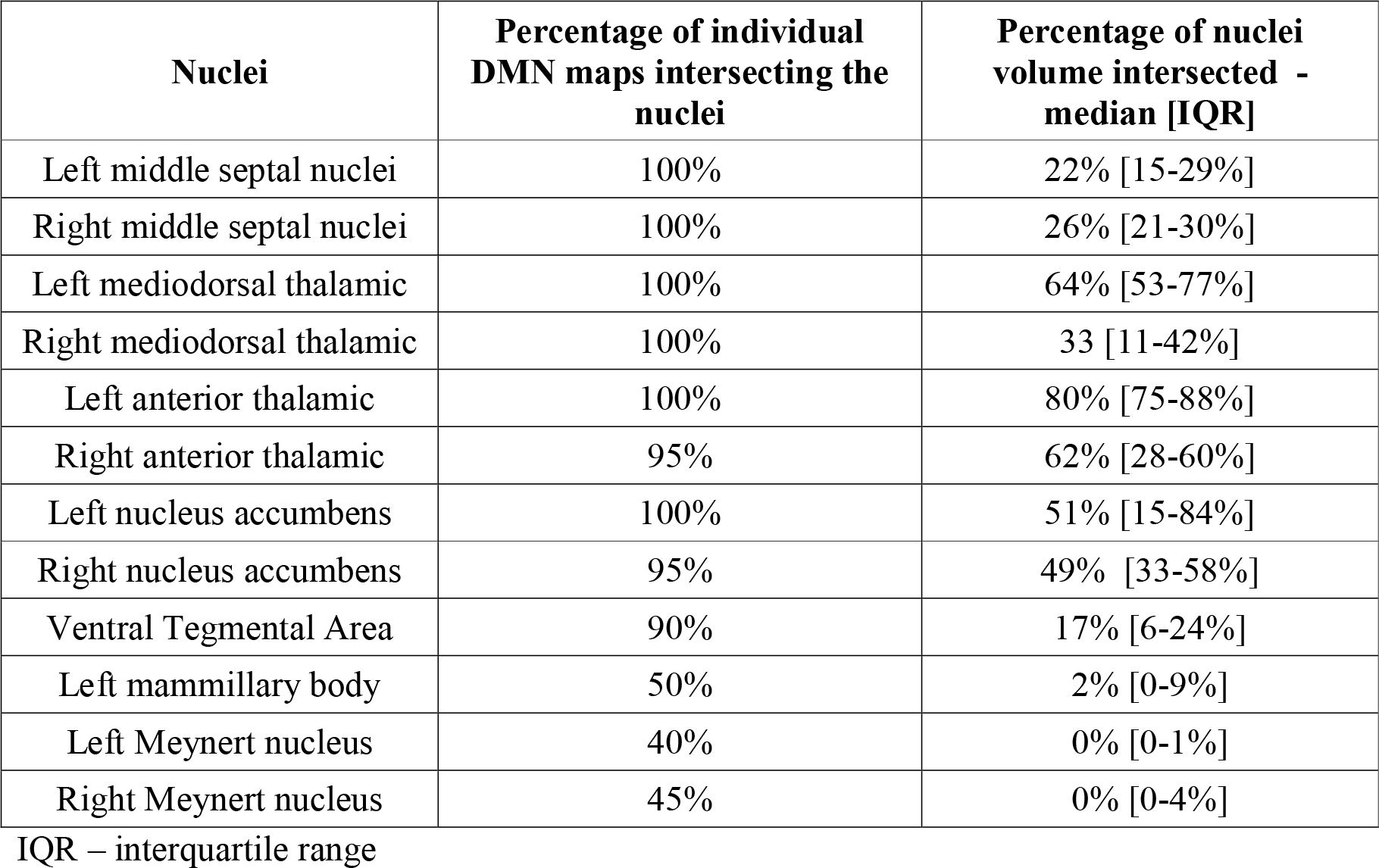
Percentage of individual DMN maps that intersect each subcortical nucleus and distribution of the volume intersected

**Figure 4.**
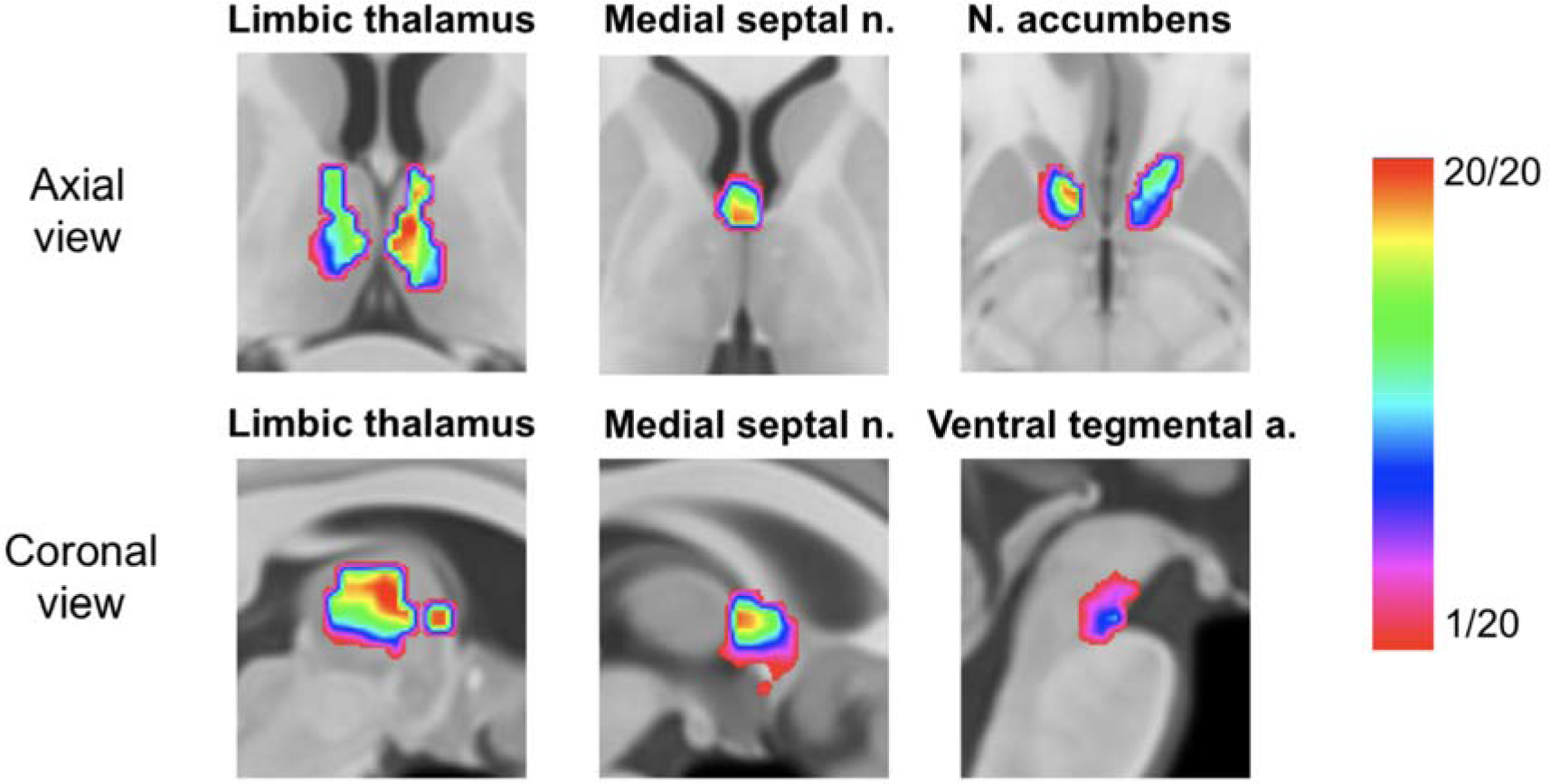
Density maps of the functionally-aligned individual DMN networks superimposed in the MNI152 space. Colour bar represents the proportion of individuals with a significant correlation in each voxel.

### 3.4 Tractography

We explored the structural connectivity of our new model of the DMN using tractography imaging techniques. Twenty-four regions of interest were defined based on the DMN we obtained in the functional space and concordant with the previous anatomical models of the DMN (Andrews-Hanna et al., 2010; Buckner et al., 2008). According to our new findings, nine additional regions of interest were defined including the left and right thalamus, the left and right basal forebrain, the midbrain and the left and right ventral prefrontal cortex (VPFC) and left and right caudate nucleus (inferior regions of the nuclei) for a total of 33 regions of interest. Figure 5 represents the structural connectivity of the network.

**Figure 5:**
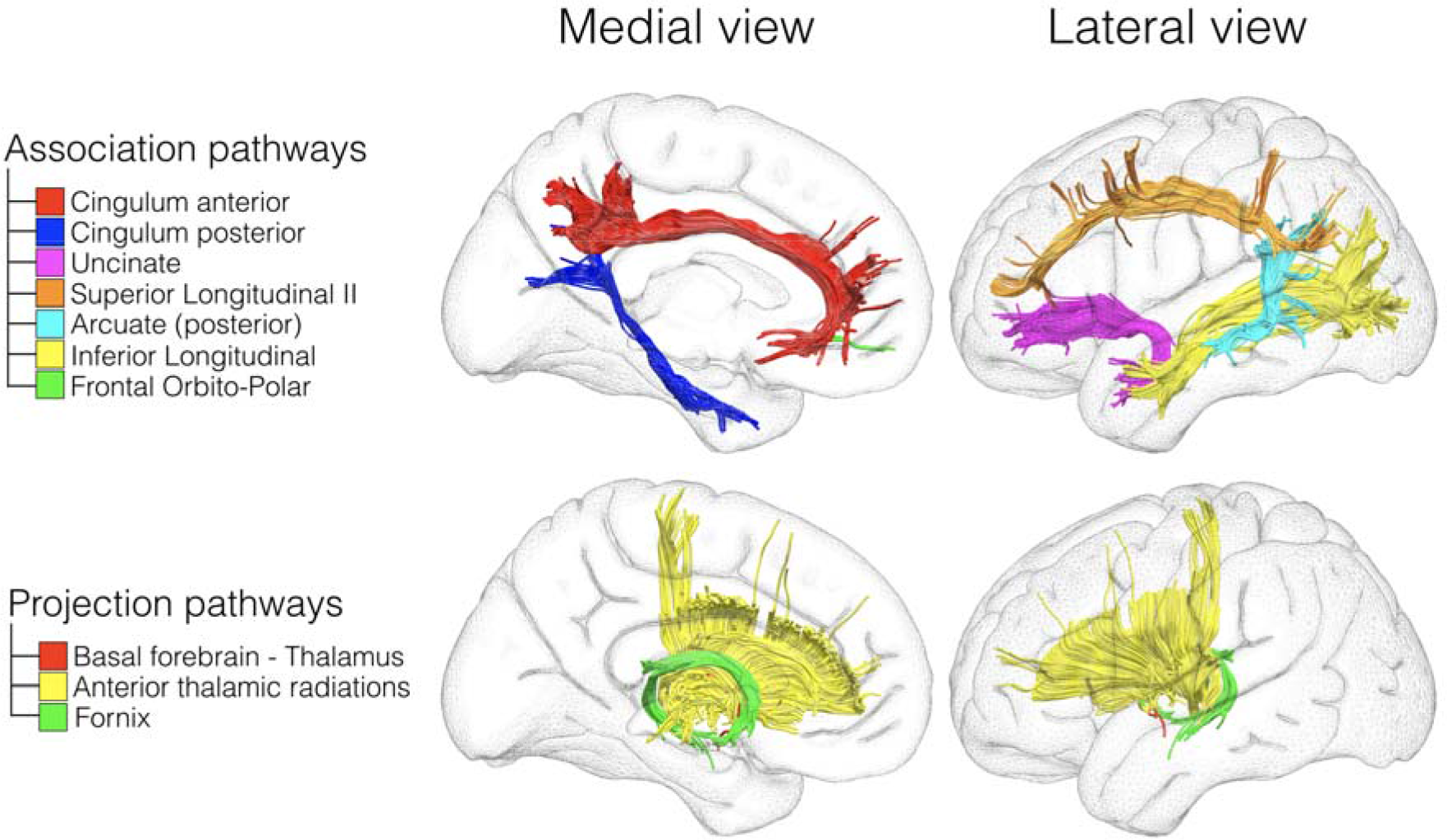
Structural connections supporting the DMN. The upper panel corresponds to the association pathways connecting the cortical regions of the DMN. The lower panel illustrates the projection pathways mediating the connections between subcortical and cortical regions of the DMN.

Results indicated that both anterior and posterior portions of the cingulum, as well as inferior longitudinal fasciculus, the second branch of the superior longitudinal fasciculus, the posterior segment of arcuate fasciculus, the uncinate fasciculus and some fibres of the frontal orbito-polar tract (Figure 5, upper panel) connected the different nodes of the DMN. Additionally, the anatomical connectivity of the basal forebrain and the thalamus with other regions of interest included: the anterior thalamic projections, connecting thalamus with medial prefrontal cortex; the cingulum, connecting basal forebrain with medial prefrontal cortex and posterior cingulate cortex; the fornix, connecting basal forebrain (specifically the region correspondent to the medial septal nuclei) to the hippocampus; and fibres connecting basal forebrain and thalamus, some of the most medial possibly corresponding to the bundle of Vicq D’Azyr (Figure 5, lower panel)

### 3.5 Graph theory analysis

Figure 6 represents the analysis of the DMN structural network with a graph theory approach using the 33 regions of interest defined from the DMN in the functional space.

Results indicate that high degrees and high betweenness centrality in the network were obtained for the basal forebrain and thalamic regions, alongside the medial prefrontal cortex and the posterior cingulate–retrosplenial cortex as well as in regions which were previously considered as hubs in the DMN.

More precisely, the [maximum-minimum] range of distribution of node degrees was [12 - 1] and the median [interquartile range] was 5 [8 - 2]. The node degrees of the left and right thalamus were 9 and 7, and of the left and right basal forebrain were 8 and 7, respectively (Supplementary material, table 1). Therefore, thalamus and basal forebrain are among the structures in the network that have connections with a high number of nodes. For betweenness centrality, the [maximum-minimum] range of distribution was [0.104 - 0] and the median [interquartile range] was 0.004 [0.03 - 0.001]. The betweenness centrality of the left and right thalamus was 0.03 and of the left and right basal forebrain were 0.03 and 0.02, respectively (Supplementary material, table 1). Hence, thalamus and basal forebrain make part of a high fraction of shortest paths in the network, that is, the shortest connections between two nodes.

**Figure 6:**
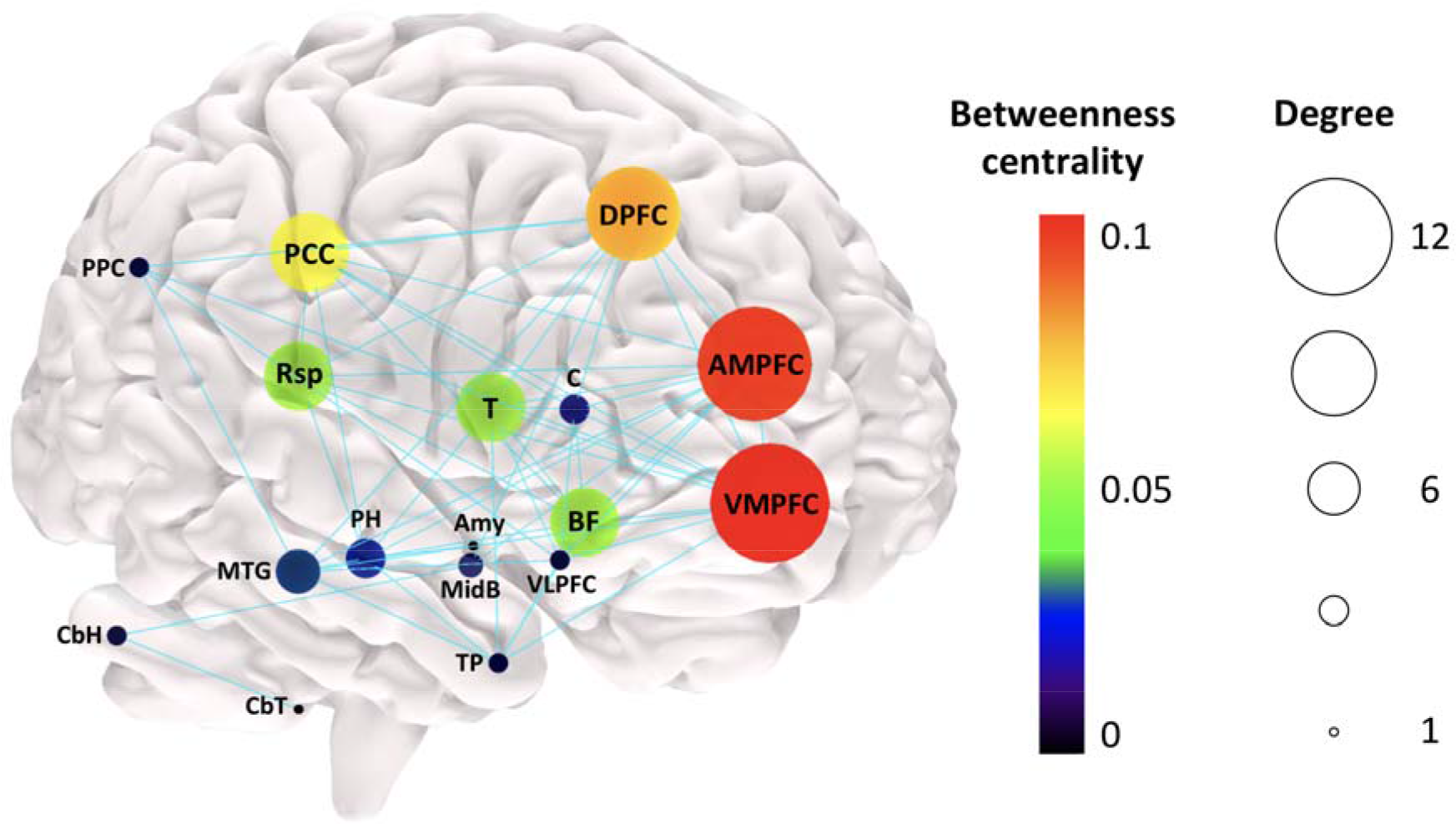
Graph theory analysis of structural connectivity. The node size represents node degree and the node colour illustrates node betweenness centrality. The edges denote presence of structural connection. DPFC, Dorsal Prefrontal Cortex; PPC, Posterior Parietal Cortex; VLPFC, Ventro-Lateral Prefrontal Cortex; Rsp, Retrosplenial Cortex; MTG, Middle Temporal Gyrus; PCC, Posterior Cingulate Cortex; C, Caudate; DPFC, Dorsal Prefrontal Cortex; AMPFC, Antero-Median Prefrontal Cortex; VMPFC, Ventro-Median Prefrontal Cortex; TP, Temporal Pole; BF, Basal Forebrain; T, Thalamus; PH, Parahippocampal Region; CbH, Cerebellar Hemisphere; CbT, Cerebellar Tonsil; Amy, Amygdala; MidB, Midbrain.

## 4. Discussion

In this study, we revisited the constituent elements of the default mode network (DMN) using an optimized method of coregistration in a functional space, besides the conventional structural alignment. Three main findings emerge from this research in healthy humans. First, higher functional connectivity correlation and sharper anatomical details were achieved when registering the DMN maps in a functional space. Second, we confirmed the hypothesis that structures of basal forebrain and anterior and mediodorsal thalamic nuclei belong to the DMN. Lastly, we characterised in detail the structural connectivity underlying functional connectivity. Based on these findings, we provided a more comprehensive neurobiological model of the DMN that bridge the gap between local differences in subcortical structures and global differences in the DMN reported in clinical studies.

The difference between alignment in the functional space and the structural space was characterised by an increase of the connectivity strength across the brain as well as in many subcortical areas classically not considered to be constituent nodes of the DMN. As previously reported, this confirmed that registration in the functional space provides a more accurate interindividual anatomical description and is recommended when doing functional connectivity analyses (Langs et al., 2015; Mueller et al., 2013).

The maps of the DMN registered in the functional space revealed previously underappreciated parts of this network such as basal forebrain and anterior and mediodorsal thalamic nuclei. Tractography analysis yielded structural connectivity of these new DMN regions to the other regions of the network. Mainly, the cingulum connected the basal forebrain with medial prefrontal cortex, posterior cingulate region, retrosplenial cortex, and hippocampus and parahippocampal regions (Catani et al., 2012, 2002). The fornix linked basal forebrain, specifically the medial septal nuclei, with the hippocampus and parahippocampal regions (Aggleton, 2000; Catani et al., 2013; Saunders and Aggleton, 2007). The anterior thalamic projections connected the thalamus with medial and ventrolateral prefrontal regions (Behrens et al., 2003) and finally some of the most medial fibres connecting the basal forebrain with the thalamus, probably corresponded to the mammillothalamic tract of Vicq D’Azyr (Balak et al., 2018; Vicq D’Azyr, 1786).

Our graph theory approach (Bullmore and Sporns, 2009) applied to the measures of structural connectivity revealed a high node degree of the basal forebrain and the thalamus in the network, as well as high betweenness centrality (Rubinov and Sporns, 2010). These results indicate that basal forebrain and thalamus have high centrality within the network, and therefore can have an important role for network integration and resilience (Bullmore and Sporns, 2009; Hagmann et al., 2008; Rubinov and Sporns, 2010), along with the classically defined hubs such as the medial prefrontal region as well as posterior cingulate and retrosplenial region (van Oort et al., 2014). Our results are concordant with recent neurophysiological evidence in rats about the influence of basal forebrain in the regulation of the DMN (Nair et al., 2018).

The involvement of basal forebrain and anterior and mediodorsal thalamus in the DMN has theoretical and functional repercussions beyond the purely anatomical level. For instance, Kernbach et al. recently demonstrated that grey-matter variability across the DMN can well predict population variation in the microstructural properties of the anterior thalamic radiation and the fornix in 10,000 people (Kernbach et al., 2018). As another example, Margulies et al. studied functional gradients along cortical surface and found that DMN areas were at the opposite end of primary motor/sensory areas in a spectrum of connectivity differentiation and that DMN areas exhibit the most considerable geodesic distance at the cortical level, being equidistant to the unimodal cortical areas (Margulies et al., 2016). These investigators suggested that the DMN acts as a neural relay for transmodal information. We speculate that thalamus and basal forebrain may follow the same model at a subcortical level, integrating functional networks related to primary functions and brainstem inputs to the associative areas (Dringenberg and Olmstead, 2003; Mease et al., 2016). The involvement of the anterior and mediodorsal thalamic nuclei as well as the basal forebrain are concordant with the role of the DMN in memory processes (Andrews-Hanna et al., 2010; Schacter et al., 2007), as all these regions are relays of the unitary model of the limbic system (Catani et al., 2013; MacLean, 1952, 1949). Previous reports of engagement of the mediodorsal thalamic nucleus and the DMN during memory tasks and the memory deficits provoked by lesions of the anterior and the mediodorsal thalamic nuclei also support this claim (Rabin et al., 2010; Spreng and Grady, 2010; Child and Benarroch, 2013; Danet et al., 2015). At a neurochemical level, the basal forebrain is also a principal actor in the production of acetylcholine (Zaborszky et al., 2015). Acetylcholine has a physiological and a neuropharmacological effect on memory processes. For instance, cholinergic system mediates rhythmic oscillation in the hippocampus that facilitates encoding (Hasselmo, 2006; Zaborszky et al., 2008). By providing evidence of the involvement of medial septal cholinergic nucleus and its structural connection to the hippocampus in our DMN model, the present work indicates a match between connectivity, neurochemistry and cognition.

The same correspondence between connectivity, neurochemistry and cognition applies to the relation between DMN and emotional modulation (Alcalá-López et al., 2017; Bzdok et al., 2013; Mars et al., 2012; Raichle, 2015; Spies et al., 2017; Zhao et al., 2017). The nucleus accumbens is a central output for the dopaminergic projections and is involved in emotion regulation and affect integration (Floresco, 2015; Laviolette, 2007). The nucleus accumbens also receives glutamatergic inputs from the hippocampus and the prefrontal cortex (Britt et al., 2012) belonging to the DMN. Surprisingly, our analysis also revealed the ventral tegmental area, which is also a dopaminergic nucleus with projections to the nucleus accumbens and the medial prefrontal cortex (Morales and Margolis, 2017). This association with the mesolimbic dopaminergic pathway is reinforced by our results of functional connectivity, since ventromedial prefrontal cortex and midbrain were among the structures with highest partial correlations with basal forebrain. Hence, integrating present and previous findings, it appears that the DMN, as defined by functional connectivity, is at the interplay between a cholinergic and a dopaminergic system dedicated to memory and emotion.

The new DMN’s subcortical structures identified in the current work have cognitive and neurochemical roles that open a new window to the understanding of distinct brain pathologies affecting DMN connectivity (Table 4). Indeed, functional connectivity in each area of the DMN is an estimate of the global coherence of the DMN. Since we demonstrated that limbic thalamus and basal forebrain are nodes with high degree and high centrality in the DMN, damage in these structures should provoke a drastic decrease of functional connectivity in the whole DMN (van den Heuvel and Sporns, 2013). For instance, Alzheimer’s disease is associated with degeneration of the cholinergic system, including the medial septal nuclei, even in the earliest clinical stages of Mild Cognitive Impairment (Grothe et al., 2010) and apparently related to significant changes of the DMN (He et al., 2007; Persson et al., 2008). The high centrality of the basal forebrain in the DMN network may explain this early link between DMN and Alzheimer’s disease. In schizophrenia, also associated with changes in the DMN connectivity (Bluhm et al., 2007; Pomarol-Clotet et al., 2008), neuropathological evidence suggests an abnormal glutamatergic-dopaminergic interaction at the level of nucleus accumbens (McCollum and Roberts, 2015). Additionally, the ventral tegmental area is connected to the nucleus accumbens through the mesolimbic system, the classical dopaminergic pathway associated with schizophrenia, and functional data shows a decrease of connectivity between VTA and several brain regions, including the thalamus, in unmedicated schizophrenic patients (Hadley et al., 2014). The pathophysiology of others diseases, such as drug addiction, depression, temporal lobe epilepsy and attention deficit and hyperactivity disorder involve modifications in the nucleus accumbens, medial septal nuclei or the thalamic nuclei that connect limbic regions as well as dysfunctional connectivity of the DMN (Table 4; Butler et al., 2013; Dinkelacker et al., 2015; Ivanov et al., 2010; Scofield et al., 2016; Vialou et al., 2010; Voets et al., 2012; Volkow et al., 2011; Yamamura et al., 2016; Zhu et al., 2016). Hence, the involvement of the basal forebrain and the thalamic nuclei in the DMN appears to bridge the gap between the subcortical anatomical differences and the global differences in the DMN previously reported.

**Table 4.** Pathophysiological associations of the subcortical structures of DMN. MCI: Mild Cognitive Impairment

| **Disease associated with DMN dysfunction** | **Pathophysiological associations with subcortical DMN** | **References** |
| --- | --- | --- |
| Alzheimer's disease | Degeneration of cholinergic system (including medial septal nuclei), even in MCI stage | Grothe et al., 2010 |
|  | Dysfunction of limbic thalamus in early stages of the disease | Aggleton et al., 2016 |
|  | Decreased functional coherence and deactivation in hub regions of DMN | He et al., 2007; Persson et al., 2008 |
| Schizophrenia | Abnormalities in glutamatergic-dopaminergic interaction in Nucleus Accumbens | McCollum and Roberts, 2015 |
|  | Decreased functional connectivity between ventral tegmental area and thalamus | Hadley et al., 2014 |
|  | Changes in the functional connectivity or activation pattern of hub regions of DMN | Bluhm et al., 2007; Pomarol-Clotet et al., 2008 |
| Temporal Lobe Epilepsy (TLE) | Dysfunctional increase in hippocampus - mediodorsal thalamus connectivity | Dinkelacker et al., 2015 |
|  | Stimulation of anterior thalamic nucleus is efficacious in treatment of TLE | Osorio et al., 2007; Salanova et al., 2015 |
|  | Enlargement of medial septal nuclei in TLE | Butler et al., 2013 |
|  | Decreased functional connectivity between hippocampus and DMN | Voets et al., 2012 |
| Depression | Resting activity of the thalamus predicts response to anti-depressant medication | Yamamura et al., 2016 |
|  | Nucleus accumbens mediates response to stress and to anti-depressant medication | Vialou et al., 2010 |

Interestingly, the thalamus and the basal forebrain are phylogenetically older than many cortical structures and especially those that compose the DMN (Butler, 2008; Karten, 2015; Yamamoto et al., 2017). The inclusion of these structures in the anatomical model of the DMN can open a window to the exploration of DMN in other mammalian species as well (Buckner and Margulies, 2018; Lu et al., 2012; Rilling et al., 2007; Vincent et al., 2007). For instance, the medial thalamus is very involved in the rat and mouse’s DMN (Gozzi and Schwartz 2016, Bertero et al. 2018).

One limitation of the present work is that the overlap between the DMN map and the nuclei studied is based on comparison with templates or variability maps, and not with the individual location of the nucleus in the explored subjects. The limited capacity of structural MRI to differentiate these small nuclei does not allow such comparison. Besides, the reverse transformation of the DMN from the functional space to the MNI space may not be exact, due to inherent limitations of inverse transformations. Although these limitations may decrease the accuracy of the intersection quantification with discrete nuclei, they did not alter the apparent overlap with basal forebrain and with the thalamus.

In conclusion, this work demonstrates that the registration of individual DMN maps in a functional space improves the definition of the anatomy of DMN by including additional structures, such as the thalamus and basal forebrain. Future research should focus on the cascade of neurochemical and pathophysiological events that follow small subcortical lesions of the DMN.

## Acknowledgement

PNA salary was supported by the Research Experience Fellowship grant of the European Academy of Neurology. The research leading to these results received funding from the “Agence Nationale de la Recherche” [grant number ANR-13-JSV4-0001-01], the European Research Council (ERC) Consolidator Grant “DISCONNECTOME” [ERC-2018-COG 818521] and from the Fondation pour la Recherche Médicale (FRM DEQ20150331725). Additional financial support comes from the program “Investissements d’avenir” ANR-10-IAIHU-06.

