## Supplementary material, table 1 for "Subcortical Anatomy of the Default Mode Network: a functional and structural connectivity study"

Table 1 – Node degree and betweenness centrality of the network nodes

| **Nodes** | **Node degree** | **Betweenness centrality** |
| --- | --- | --- |
| Right Ventro-Medial Prefrontal Cortex | 12 | 0,104 |
| Left Ventro-Medial Prefrontal Cortex | 12 | 0,070 |
| Right Antero-Medial Prefrontal Cortex | 12 | 0,060 |
| Left Antero-Medial Prefrontal Cortex | 11 | 0,040 |
| Left Dorsal Prefrontal Cortex | 10 | 0,075 |
| Right Dorsal Prefrontal Cortex | 10 | 0,057 |
| Left Posterior Cingulate Cortex | 9 | 0,025 |
| Left Thalamus | 9 | 0,029 |
| Left Basal Forebrain | 8 | 0,032 |
| Right Posterior Cingulate Cortex | 8 | 0,028 |
| Left Retrosplenial Cortex | 8 | 0,020 |
| Right Basal Forebrain | 7 | 0,024 |
| Right Retrosplenial Cortex | 7 | 0,020 |
| Right Thalamus | 7 | 0,033 |
| Left Caudate | 5 | 0,003 |
| Left Parahippocampal region | 5 | 0,003 |
| Right Middle Temporal Gyrus | 5 | 0,020 |
| Left Middle Temporal Gyrus | 4 | 0,013 |
| Left Ventral Prefrontal Cortex | 4 | 0,002 |
| Right Parahippocampal region | 4 | 0,004 |
| Left Temporal Pole | 3 | 0,002 |
| Right Caudate | 3 | 0,001 |
| Left Posterior Parietal Cortex | 3 | 0,002 |
| Midbrain | 3 | 0,000 |
| Right Cerebellar hemisphere | 2 | 0,000 |
| Right Temporal Pole | 2 | 0,003 |
| Right Posterior Parietal Cortex | 2 | 0,001 |
| Left Ventral Prefrontal Cortex | 2 | 0,000 |
| Left Amygdala | 1 | 0,000 |
| Left Cerebellar Hemisphere | 1 | 0,000 |
| Left Tonsil | 1 | 0,000 |
| Right Amygdala | 1 | 0,000 |
| Right Tonsil | 1 | 0,000 |
